# The possible fidelity-speed-proofreading cost trade-offs in DNA replication due to the exonuclease proofreading

**DOI:** 10.1101/2021.02.18.431768

**Authors:** Qiu-Shi Li, Yao-Gen Shu, Wen-Bo Fu, Zhong-Can Ou-Yang, Ming Li

## Abstract

DNA replication is a high-fidelity information-copying processes which is realized by DNA polymerase (DNAP). The high fidelity was explained on the basis of the well-known kinetic-proofreading mechanism (KPR), under which the so-called fidelity-speed trade-off was studied theoretically. However, numerous biochemical experiments have shown that the high fidelity of DNA replication is achieved due to the initial discrimination of polymerase domain of DNAP, as well as the proofreading of the exonuclease domain of DNAP. This exonuclease-proofreading mechanism (EPR) is totally different from KPR. So the trade-off issues are worth being re-examined under EPR. In this paper, we use the first-passage method recently proposed by us to discuss the possible trade-offs in DNA replication under EPR. We show that there could be no fidelity-speed trade-off under EPR, i.e., the fidelity and the speed can be simultaneously enhanced by EPR in a large range of kinetic parameters. This provides a new perspective to understand the experimental data of the exonuclease activity of T7 DNAP and T4 DNAP. We also show that there exists the fidelity-proofreading cost trade-off, i.e., the fidelity is enhanced at the cost of increasing the futile hydrolysis of dNTP. A possible way to avoid this trade-off is to regulate the rate of DNAP translocation: slowing down the forward translocation (in the presence of the terminal mismatch) can enhance the fidelity without changing the speed and the proofreading cost. Our theoretical analysis offers deeper insights on the kinetics-function relation of DNAP.

PACS numbers: 82.39.-k, 87.15.Rn, 87.16.A-

## I. INTRODUCTION

High fidelity is vital to biosynthetic processes. For example, the fidelity can be as high as ~ 10^4∼10^ for DNA replication[1], 10^4^ for transcription, and ~ 10^3∼4^ for translation[2]. In 1970s’, the kinetic proofreading mechanism (KPR) was proposed by Hopfield[3] and Ninio[4] to explain such high fidelity. The Hopfield model is conceptually shown in Fig.1(A1), in which the biosynthetic reaction is regarded as a binary copolymerization process of Right(R) and Wrong(W) substrates. The fidelity is determined by the huge difference between the incorporation rate of R and that of W. However, numerous experiments of structural biology and biochemistry have shown that KPR is suitable for translation but not for DNA replication and RNA transcription. DNA replication is catalyzed by DNA polymerase (DNAP). DNAP has two domains, the polymerase domain (Pol) which can incorporate nucleotides (dNTP) and the exonuclease domain (Exo) which can excise the just-incorporated nucleotide (NMP). The exonuclease proofreading mechanism (EPR) is schematically shown in Fig.1(B1) which is totally different from the KPR. The core assumption of KPR is that the proofreading occurs before the dNTP incorporation and thus the reaction pathway of incorporating either R or W can be essentially regarded as irreversible (Fig.1(A2)). For such schemes, simple mathematical treatments like Michaelis-Menten kinetics can be used to calculate the fidelity and other quantities. For the EPR, however, the proofreading occurs after dNTP incorporation, i.e., the terminal NMP of the primer can only be excised after the terminal is transferred to Exo. So the reaction scheme under EPR can no longer be regarded as irreversible. Instead, it has to be regarded as apparently reversible like Fig.1(B2). For such schemes, Michaelis-Menten kinetics is invalid, and one should use the copolymerization theory to analyze the polymerization process, such as the steady-state copolymerization theory[5, 6], the first-passage (FP) method[7] developed by us and the iterated function system (IFS) developed by P.Gaspard[8].

**FIG. 1:**
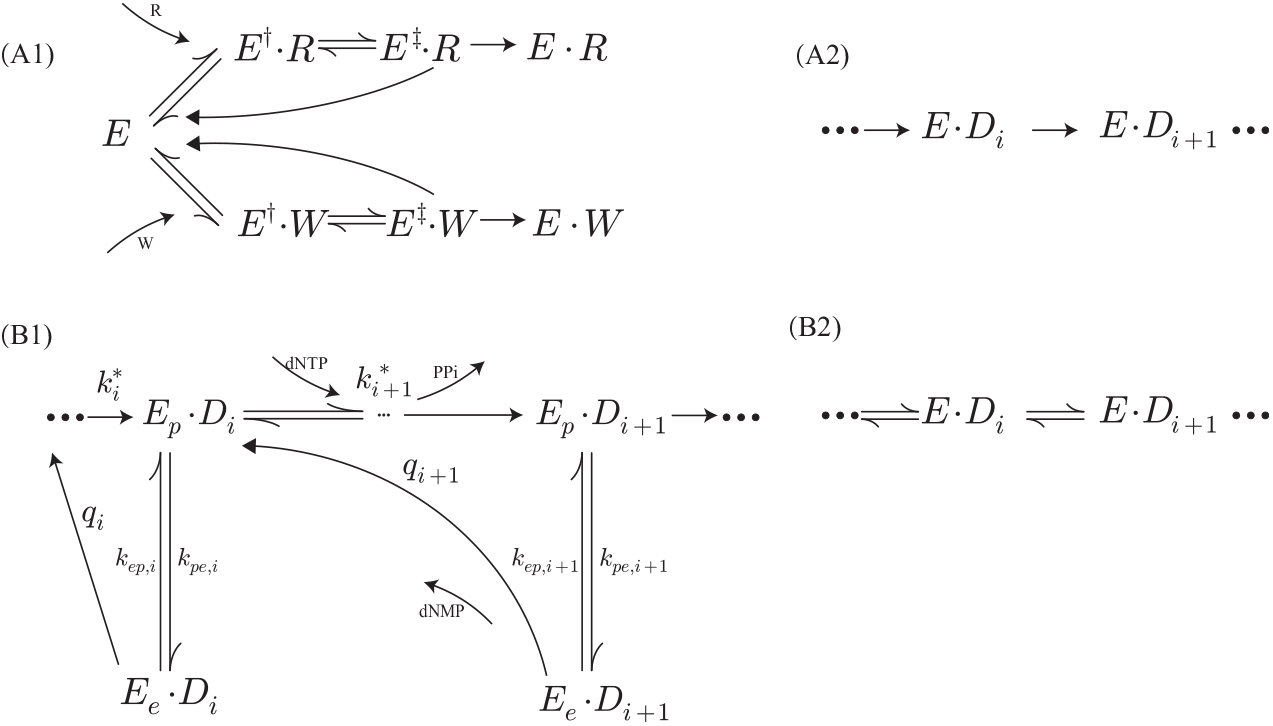
(A1)The kinetic proofreading model. (B1) The real reaction scheme of DNA replication. *E_p_* and *E_e_* indicate respectively that the primer terminal is at Pol or Exo. *D_i_* represents the primer of length *i*. *k*\* represents the effective incorporation rate. *k_pe_* and *k_ep_* represent the transfer rates between Pol and Exo. *q* represents the excision rate. When only calculating the replication fidelity, either pathway in (A1) can be reduced to (A2), and (B1) can be reduced to (B2).

It is widely believed that the replication fidelity can only be enhanced at the cost of decreasing the speed, because of the proofreading (the so-called fidelity-speed trade-off). Previous theoretical studies under KPR show that there may be the fidelity-speed trade-off, e.g. in protein synthesis[9, 10]. Some work studied the generalized proofreading network and found a regime that the speed can be highly increased with the fidelity little decreased[11] which may exist in the charging of tRNA by tRNA synthetases, RecA filament assembly on ssDNA, and protein synthesis. Nevertheless, as mentioned above, the EPR in the DNA replication is totaly different from KPR, so all the conclusions obtained under KPR are worthy to be re-examined under EPR. In this paper, we try to investigate the possible trade-offs in DNAP replication under EPR. The contribution of the exonuclease to the overall fidelity has been discussed in details in Ref.[12] using the FP method. Other important quantities like the replication speed and the proofreading cost will be discussed in this paper. The rest of the this paper is organized as following. A brief review of the FP method and the overall fidelity will be introduced first. Then the replication speed and the proofreading cost will be defined and calculated by the FP method. We will try to clarify the possible trade-offs between the fidelity, the speed and the proofreading cost. Finally we will discuss the DNAP translocation which may have a substantial impact on the above trade-offs.

## II. RESULTS AND DISCUSSION

### A. There can be no fidelity-speed trade-off due to the exonuclease proofreading

Consider the replication process shown in Fig.1(B1) along a template sequence *X*_1_*…X_i_…X_L_*. The primer grows from *α*_1_ until reaches *α*_1_*…α_L_*. *X* and *α* can be any one of the four dNTP (*A, T, G, C*). In Fig.1(B1), there are indeed multiple steps to incorporate dNTP, as indicated by *k*\*. For brevity to illustrate the basic logic, however, we only discuss a hypothetic case in which the dNTP incorporation is a one-step process with rate constant *k*\*. The major conclusions given below still hold for the multi-step dNTP incorporation process.

Let 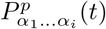 and 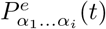 represent the probability of the intermediate product of the sequence *α*_1_*…α_i_* at time *t* with the primer terminal in Pol or Exo respectively. The time-dependent probabilities can be described by the following master equations,

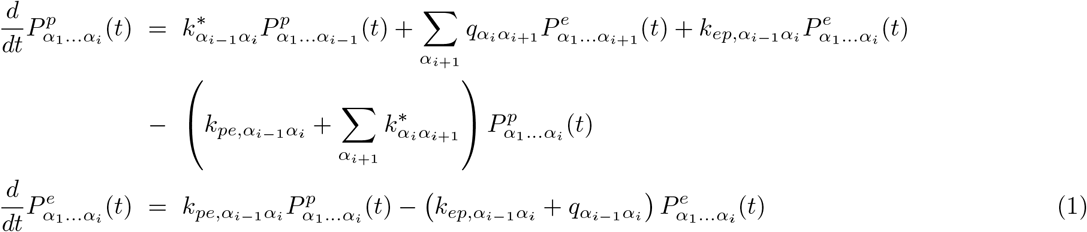

In this paper we only consider the first-order neighbor effect (as indicated by the subscripts *α_i_*_−1_*α_i_*). 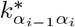 represents the rate of incorporating *α_i_* to the terminal 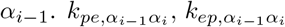 and 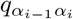 correspond to *k_pe,i_*,*k_ep,i_ q_i_* in Fig.1(B1) with the terminal sequence *α_i_*_−1_*α_i_* respectively. To simplify the notation, we have neglected the superscript *X*_1_*…X_i_…X_L_* in all the kinetic parameters.

To calculate the local fidelity at template site *i*, we need to calculate the probability distribution of all the final products 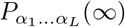. We define the probability to find *α_i_* at site *i* as 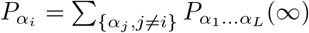 and then define the fidelity at site *i* as 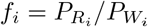 for any particular mismatch *W_i_*. By using the FP method, we get the approximate analytical expression of the local fidelity in a vast range of rate parameters,

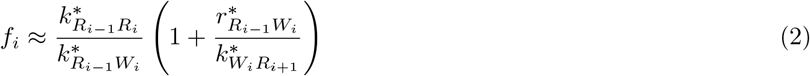

Here *r*\* = *k_pe_q/*(*q* + *k_ep_*). Detailed information can be found in Appendix A and Ref.[7]. To simplify the discussion below but not to lose generality, we consider just one type of mismatch *W_i_*.

Let *r*\* = 0 in Eq.(2) (*i.e. k_ep_* = 0), one can get the fidelity without exonuclease proofreading,

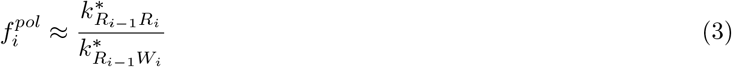

Comparing Eq.(2) with Eq.(3), we can see that the contribution of the exonuclease proofreading to the overall fidelity is proportional to 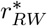.

Below we discuss the overall replication speed at site *i*. Since 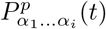 represents the probability to find the primer terminal at Pol with sequence *α*_1_*…α_i_* at time *t*,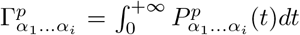 represents the residence time of the primer in this state during the entire replication process, according to the FP method (for details see Ref.[7]). This defines the total residence time at site *i*,

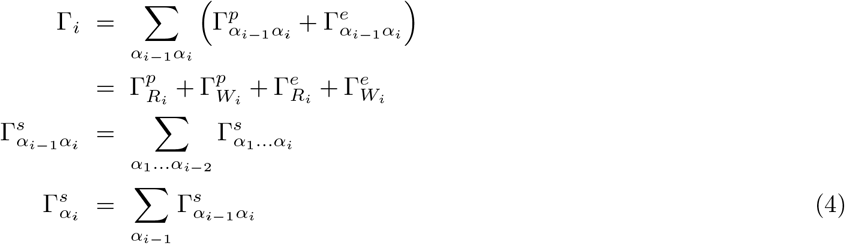

Here *s* = *p, e*. So the overall speed at site *i* can be defined as *v_i_* = 1*/*Γ_*i*_. By using the FP method, one can get

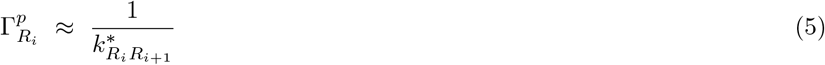

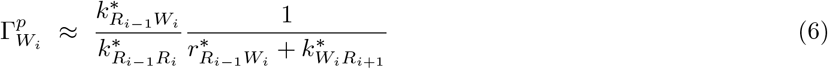

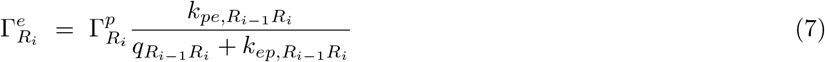

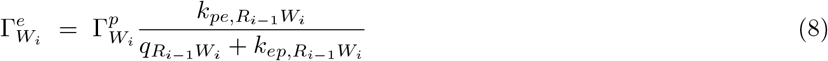

It should be noted that although all the above results are derived for the hypothetic one-step dNTP incorporation process with the rate constant *k*\*, they still hold for the real multi-step dNTP incorporation process if *k*\* is correctly understood as the effective incorporation rate which can be defined rigorously and uniquely by the FP method (for details see appendix A).

Let *r*\* = 0 in Eq.(6) and *k_pe_* = 0 in Eq.(7,8), the total residence time without exonuclease can be obtained. 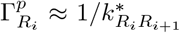 and 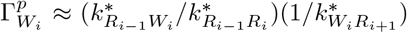 correspond to the residence time with terminal R or W respectively. Although the probability to incorporate W at site *i* is low 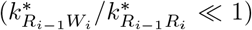, it may take much longer time 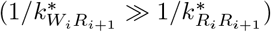 to bury the mismatch. So 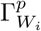 may be larger than 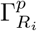. For example, the kinetic assays which were widely used to measure the effective rate of the multi-step dNTP incorporation process have shown 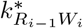 may be smaller or larger than 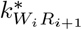 for human mitochondrial DNAP, depending on the template sequence around *i* [13] (for more detailed explanation, see Ref.[12]). For cases that 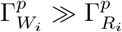, the exonuclease proofreading may largely decrease 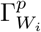 by excising rather than burring the mismatch (noting that 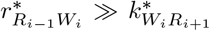) so as to enhance the overall speed significantly. For cases that 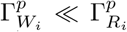, the exonuclease proofreading almost does not affect the overall speed (with only negligible enhancement).

On the other hand, the excision itself does spend extra time 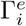. It can be clearly seen from Eq.(7,8) that 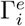 is negligible when compared with 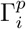, if *q* + *k_ep_ > k_pe_*. We note that some real DNAPs do meet this condition. For example, *q_RW_* = 896*s*^−1^, *k_ep,RW_* = 714*s*^−1^, *k_pe,RW_* = 2.3*s*^−1^ for T7 DNAP[14], and *q_RW_* ≈ 200*s*^−1^, *k_ep,RW_* = 526*s*^−1^, *k_pe,RW_* = 11*s*^−1^ for T4 DNAP[15]. Hence, in such a regime, the exonuclease can enhance the fidelity and the speed simultaneously, which implies at some extent the evolutionary optimization of the exonuclease activity. Whether this is true for other DNAPs remains to be checked by experiments.

It should be pointed out that such no-trade-off between the fidelity and the speed has been reported for DNA replication[16]. In that work, however, the authors did not handle the complete EPR mechanism (Fig.1(B1)) but only treated part of the reaction scheme. Moreover, they obtained the conclusions under some regime of parameters which is quite different from that discussed here. Whether such no-trade-off between the fidelity and the speed exist for a vast range of kinetic parameters is open for future studies.

### B. There does exist fidelity-proofreading cost trade-off due to the exonuclease proofreading

The enhancement of the fidelity due to exonuclease proofreading occurs definitely at the cost of increasing the amount of futile hydrolysis of dNTP. A. R. Fersht and K. A. Johnson had discussed this type of proofreading cost[14, 17], by defining the cost as the probability of dRTP being futilely hydrolyzed. K. Banerjee et al. also proposed that the fidelity-proofreading cost exist for the real DNA replication process[16]. Here we similarly define the proofreading cost as the amount of all NMP excised during the proofreading process and calculate it by the FP method. Noting that 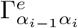 represents the total residence time that the primer terminal spends at Exo. Once the terminal is transferred to Exo, it will take the mean time 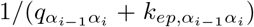 to leave this state and be excised with the probability 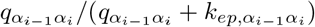. Therefore, the proofreading cost at site *i* is,

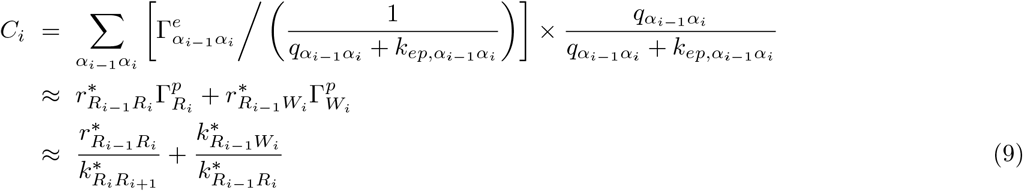

The first term 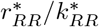 represents the amount of excised RMP. For T7 DNAP, 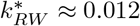[12, 18], 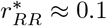[12, 14]. So the proofreading cost is almost proportional to 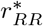.

On the other hand, the fidelity is proportional to 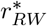. Also noting that *r*\* = *k_pe_q/*(*q* + *k_ep_*). Since the distance between the Pol and Exo is as large as 3 ~ 4*nm*[1], *q* and *k_ep_* are only determined by the Exo domain, *i.e. q_RR_ ≈ q_RW_* and *k_ep,RR_ ≈ k_ep,RW_*. So 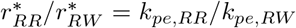. Since *k_pe,RR_* and *k_pe,RW_* are both determined by the melting propensity of the DNA duplex due to its interaction with the Pol domain, it is conceivable that *k_pe,RR_* is positively correlated with *k_pe,RW_*. So 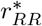 also roughly increases with increasing 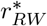. Hence, there exists the trade-off between the fidelity and the proofreading cost.

A similar conclusion was obtained in Ref.[16]. Although their treatment of the EPR mechanism is different from ours presented here, they both indicate that the proofreading cost may impede further enhancement of the fidelity for the real DNA replication.

### C. The possible way to avoid the fidelity-proofreading cost trade-off

For the real replication process, the DNAP translocation along the template must be considered, as shown in Fig.2 (details see Ref.[12]). So far as we know, there were no studies on the trade-off issues which considered the translocation step. The impact of DNAP translocation on the fidelity was recently studied by us[12]. Below we discuss how DNAP translocation affects the trade-offs between the fidelity, the speed and the proofreading cost.

**FIG. 2:**
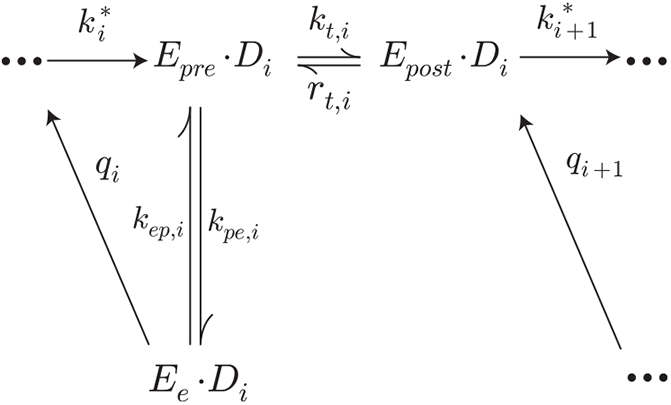
The reaction scheme considering DNAP translocation. *k_t_, r_t_* represent the forward and backward translocation rate.

Intuitively, once Pol incorporates a dWTP, DNAP may translocate forward or the terminal may be transferred to Exo and be excised there. If the forward translocation is relatively slow, it is more likely that the mismatch will be excised. According to the FP method, the fidelity now can be written as

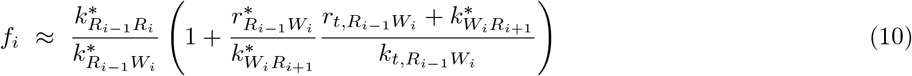

It can be seen that DNAP translocation has no impacts on the fidelity if the exonuclease is absent (*r*\* = 0), and the overall fidelity increases with decreasing 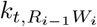.

One can also calculate the residence time

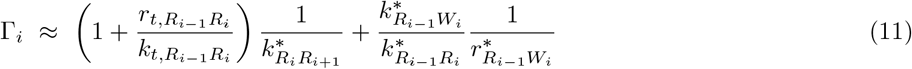

and the proofreading cost

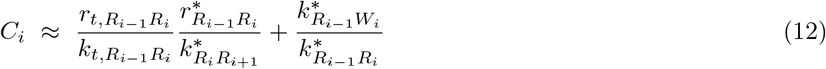

Since 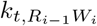 is absent from Eq.(11,12), it does not affect the speed and proofreading cost. So DNAP can possibly enhance the fidelity by decreasing 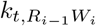 while avoid decreasing the speed or increasing the proofreading cost.

## III. SUMMARY

In this paper, we introduced the EPR mechanism which is different from the well-known KPR mechanism, and discussed the possible trade-offs between the replication fidelity, the replication speed and the proofreading cost, in the theoretical framework of EPR. We concluded that the exonuclease proofreading can enhance the fidelity and the speed simultaneously, or enhance the fidelity without decreasing the speed, by controlling some key kinetic parameters. That is to say there can be no fidelity-speed trade-off in some regime of kinetic parameters (i.e., the excision rate *q* or the transfer rate from Exo to Pol *k_ep_* is much faster than the transfer rate from Pol to Exo *k_pe_*). This condition seems to be met by some real DNAPs, indicating there may be some evolutionary optimization on the exonuclease kinetics. Hence, the discussion given here offered a new perspective to understand the design principle of DNAP.

We also showed there does exist fidelity-proofreading cost trade-off, i.e., the exonuclease can not enhance the fidelity while decreasing the proofreading cost. We propose a possible way to avoid this trade-off. If the forward DNAP translocation is quite slow when the primer terminal is a mismatch, the fidelity can possibly be enhanced without changing the proofreading cost. Unfortunately, so far there were no experiments investigating DNAP translocation with terminal mismatch. We hope the theoretical analysis presented here can attract more attention of experimentalists to examine the DNAP translocation kinetics in much more details.

Lastly, the real reaction scheme of DNA replication may be far more complex than the model we discussed in this paper. For example, the DNAP may dissociate from the DNA, the terminal transfer process may contain more sub-steps, and there could be higher-order neighbor effects. The fidelity for such complex models has been investigated in details by us[7, 12], but the speed and the proofreading cost has not been discussed ever. The theoretical analysis in this paper is hopefully generalized to study the possible trade-offs in these complex models, which will be presented elsewhere in the future.

## IV. ACKNOWLEDGMENTS

The authors thank the financial support by National Natural Science Foundation of China (No.11675180,11774358), the CAS Strategic Priority Research Program (No.XDA17010504), Key Research Program of Frontier Sciences of CAS (No.Y7Y1472Y61), Research Fund of Wenzhou Institute CAS (No.WIUCASYJ2020004,WIUCASQD2020009).

## Appendices

## APPENDIX A: BASICS OF FP METHOD

The complete master equations are,

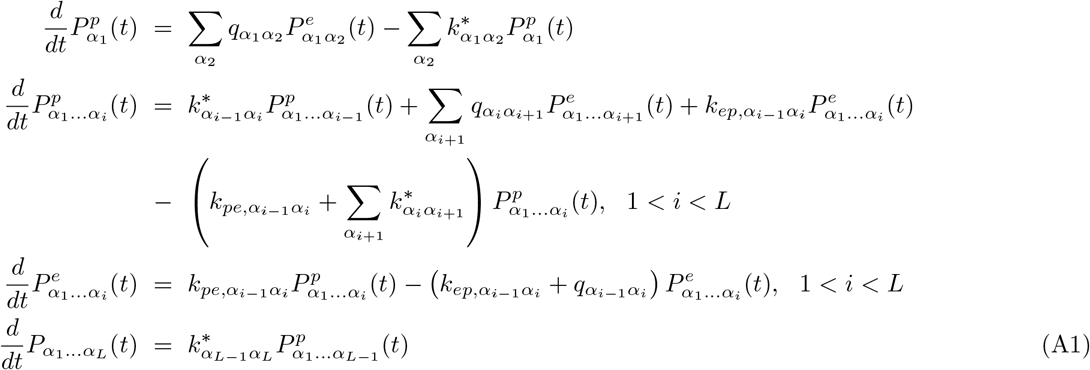

To get the probability of the final product, one can integrate Eq.(A1),

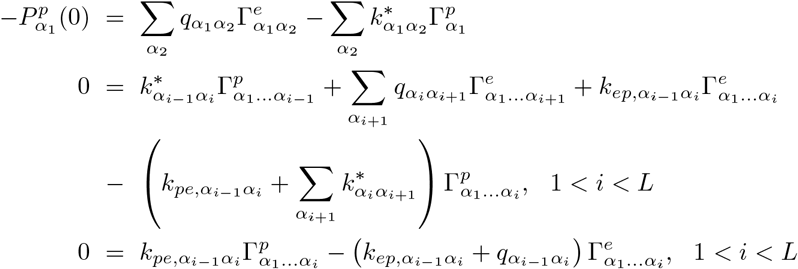

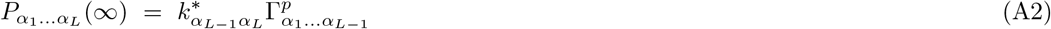

Here 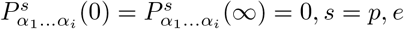 for 1 < *i* < *L*. 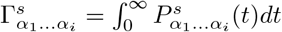. Eq.(A2) can be rewritten as

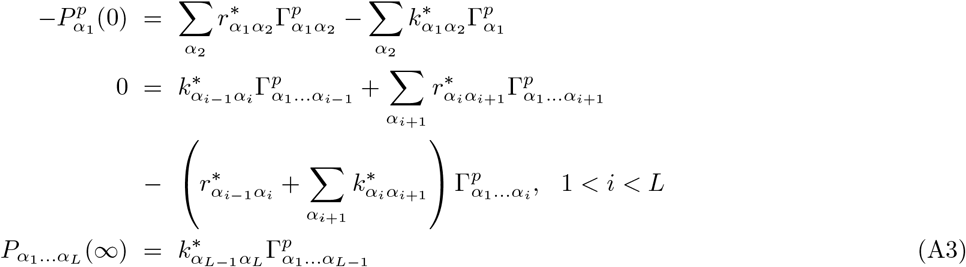

and,

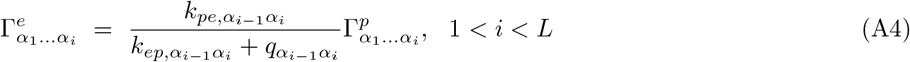

In Eq.(A3), *r*\* = *k_pe_q/*(*q* + *k_ep_*). By solving Eq.(A3), one can get (details see Ref.[7]),

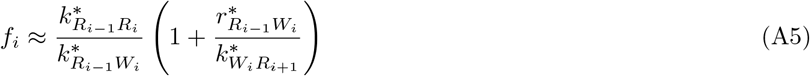

and

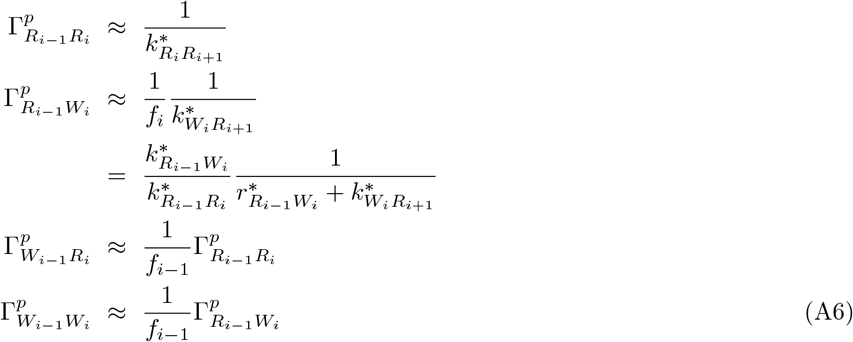

When considering the multi-step dNTP incorporation process, the original master equation is different from Eq.(A1), but the integrate master equations Eq.(A3) still holds with the uniquely defined effective rate *k*\*. An example can be found in Ref.[12] Sec III A. It should be pointed out that the residence time of all the intermediate states should be explicitly calculated (like Eq.(A4)) and be summed to get the total residence time at site *i*.

## APPENDIX B: REACTION SCHEME CONSIDERING DNAP TRANSLOCATION

For the reaction scheme considering DNAP translocation, the integrated master equations are

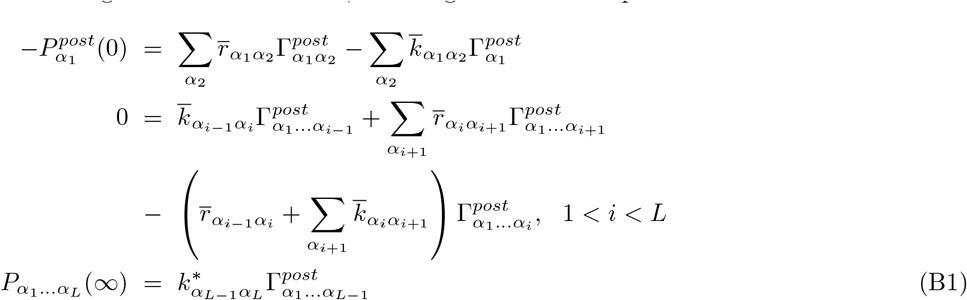

and

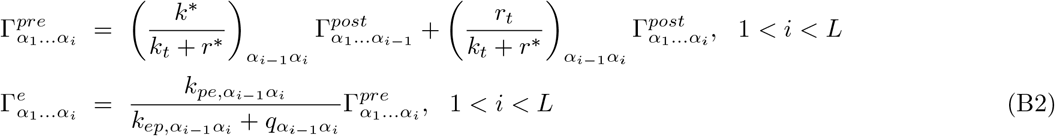

By solving Eq.(B1), one can get the fidelity

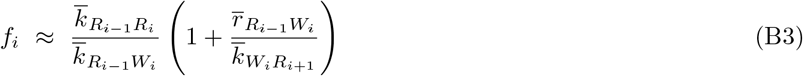

Here 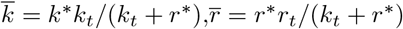.

Since DNAP translocation is always fast, *i.e.* 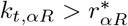, if the primer terminal is RMP, one gets 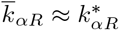, and Eq.(B3) can be rewritten as

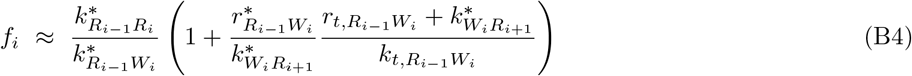

The total residence time at site *i* is 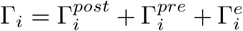. Here 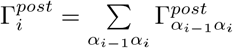

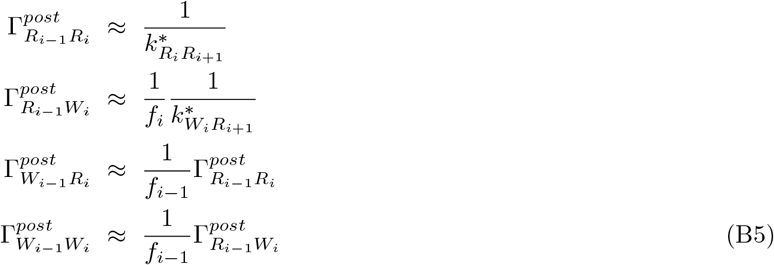

Because *f_i_*_−1_ is very large, 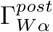 is negligible compared to 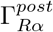.

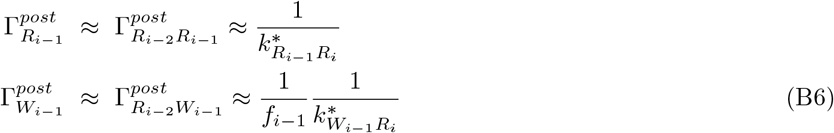

According to Eq.(B2), one can get the following to calculate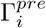

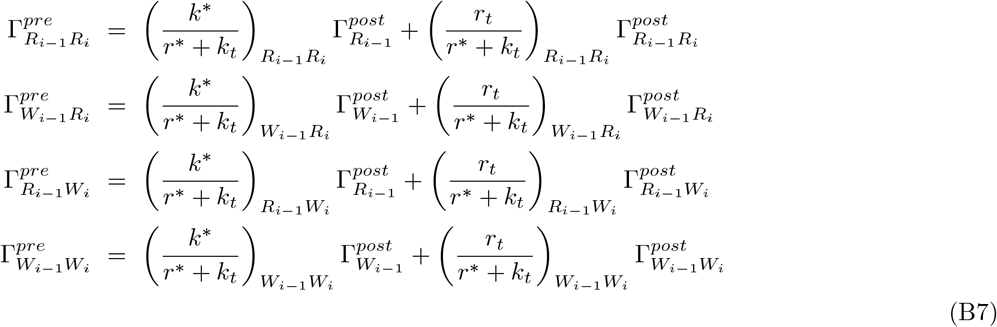

Since DNAP translocation is relative fast, *i.e.* 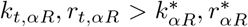, if the primer terminal is RMP, it can be seen 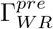 is negligible compared with 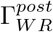 and 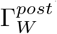. And 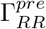 is,

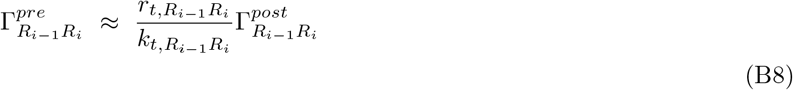

Because 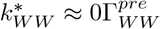 is negligible. According to Eq.(B5,B6),

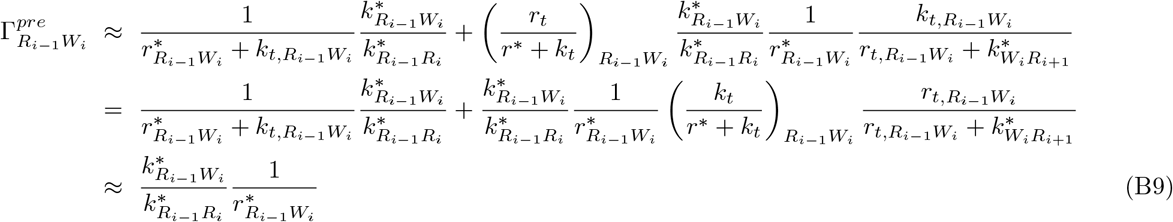

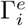 is negligible if *q* + *k_ep_ > k_pe_*, according to Eq.(B2). So we get the total residence time,

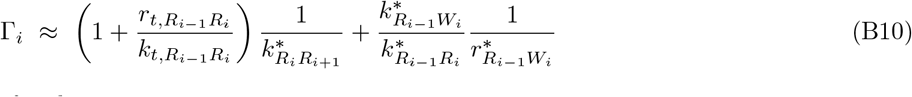

Follow the same logic, the proofreading cost is,

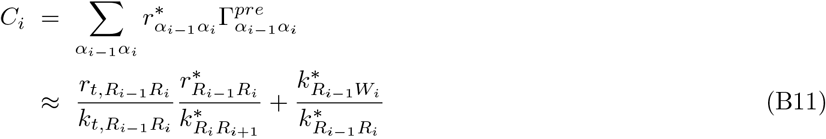

